# *VariantSpark*, A *Random Forest* Machine Learning Implementation for Ultra High Dimensional Data

**DOI:** 10.1101/702902

**Authors:** Arash Bayat, Piotr Szul, Aidan R. O’Brien, Robert Dunne, Oscar J. Luo, Yatish Jain, Brendan Hosking, Denis C. Bauer

## Abstract

The demands on machine learning methods to cater for ultra high dimensional datasets, datasets with millions of features, have been increasing in domains like life sciences and the Internet of Things (IoT). While *Random Forests* are suitable for “wide” datasets, current implementations such as *Google’s PLANET* lack the ability to scale to such dimensions. Recent improvements by *Yggdrasil* begin to address these limitations but do not extend to *Random Forest*. This paper introduces *CursedForest*, a novel *Random Forest* implementation on top of *Apache Spark* and part of the *VariantSpark* platform, which parallelises processing of all nodes over the entire forest. *CursedForest* is 9 and up to 89 times faster than *Google’s PLANET* and *Yggdrasil*, respectively, and is the first method capable of scaling to millions of features.

## 1 Introduction

The ongoing digital revolution is causing a dramatic increase in data collected about almost every aspect of life [12]. These datasets are not only growing by capturing more events (samples) but also by capturing more information about these events (features). The challenge of “big” and “wide” data is especially pronounced in the biomedical space where, for example, whole genome sequencing technology enables researchers to extract over 3 billion features from the human genome for analysis [13]. Other domains are also seeing a rapid increase in the number of features processed by statistical or machine learning applications [1].

While statistical linear models can deal with such wide datasets by analyzing each feature independently [7], there is a growing demand for more realistic approaches that can discover interacting features using machine learning [10]. In the life-science space, this would allow modeling the interactions between genes that result in complex traits like height [9] or diseases like obesity [11]. In particular, *Decision Tree* [10] based models have been successfully applied to uncover interactions between features [18].

As *Decision Trees* fitting algorithms are greedy and may yield an estimate with a high variance, Breiman developed an ensemble approach for improving accuracy by aggregating over a large number of *Decision Trees*, called *Random Forest* [5]. *Random Forest* models are particularly well suited for datasets that are wide and there is a need to capture interactions for two reasons:

- Wide datasets, particularly when there are more features than samples, cause other machine learning methods to overfit easily, whereas *Random Forest* models are more resistant to this “curse of dimensionality” [2, 4]
- There is significant scope for parallelization in *Random Forest* algorithms allowing forests to be grown efficiently even on large datasets.

The first implementations of *Random Forest* in *R* (*randomForest*) were based on the original *Fortran* code by Breiman [5]. Later *ranger* [17] provided a *C++* implementation of *Random Forest* with an *R* interface, which also covers the loading and pre-processing. *H2O* [8] provides another *R* interfacing implementation with *Scala* back end. In terms of parallelization, all of these implementations are optimized for a *high performance computing* (*HPC*) machine (a single computing node). However, the *Random Forest* algorithm allows for parallelization on a distributed computing platform.

*Apache Spark* is particularly suitable for such massively parallel interconnected calculations as it offers a distributed computing architecture that enables communication beyond compute-node boundaries in a standardized approach [13]. In the *Spark* cluster, there are several *Workers* and a *Master* each of which is a computing node in the network. The *Driver* program (run on the *Master* node) coordinates the job flow by controlling the *Executors* (run on the *Worker* nodes).

*Google’s PLANET* [3] is a *MapReduce* implementation of *Random Forest* and the first to parallelize processing each node of a tree in a distributed fashion. Hence, the ideas from *Google’s PLANET* are now used in many “Big Data” machine learning libraries, such as *Spark MLlib* and *XGBoost* [6]. *Google’s PLANET* partitions data by samples with *Workers* holding all features for a fraction of samples. However, this solution produces an approximate split and limits the depth of tree for high dimensional datasets [1].

*Yggdrasil* [1] overcomes this limitation by flipping the dataset and partitioning it by features rather than samples. The *Driver* aggregates the best local splits computed on the *Executors*, identifies the best global split, and updates the *Executors* accordingly. However, the work is limited to *Decision Trees* and does not implement bootstrapping or *mtry* (number of features considered at each node), which are essential components in a *Random Forest*.

*CursedForest* extends *Yggdrasil*’s approach to Decision Trees to *Random Forest* models. *CursedForest* also introduces a novel method of parallelization in the tree growing process such that nodes of different trees are processed in parallel. This enables highly accurate multivariate models to be built on large datasets with millions of features. For more details see Section 2.3.

In this paper we evaluated different implementations of *Random Forest*, including *Google’s PLANET*, for their ability to scale to large datasets (Section 3.1). Section 3.2 and Section 3.3 tests the limits of *CursedForest* and Section 3.4 benchmarks *CursedForest* against *Yggdrasil*. Details of *CursedForest* are elaborated in Section 2.3.

## 2 Methods

### 2.1 Datasets

For the evaluation, well-known genomic datasets (1000 Genomes Project [14]) along with a synthetic dataset are used. The genomic dataset is subsetted to create 5 datasets of different size (see Table 2). Each dataset includes the genomic profile as well as the ethnicity of a few thousand individual humans. In our evaluation, the person’s ethnicity (response variable) is predicted from his or her genomic profile and evaluated against the known ethnicity. The genomic profile is a set of features taking the values 0/0, 0/1 or 1/1 which we encode as 0, 1 and 2 respectively.

**Table 1:**
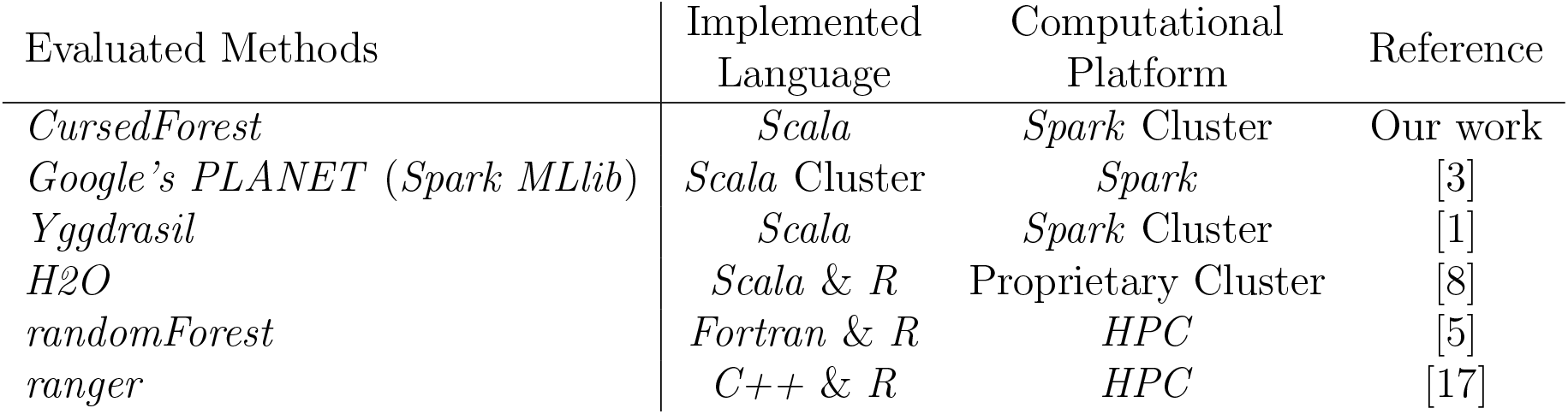
Methods evaluated

**Table 2:** Datasets used for evaluations

| Dataset | Description | Samples | Features |
| --- | --- | --- | --- |
| 80M | 1000 Genomes Phase 3 chr1 to chr22 | 2,540 | 81,047,467 |
| 20M | 1000 Genomes Phase 3 chr1 to chr3 | 2,504 | 19,328,051 |
| 6M | 1000 Genomes Phase 3 chr1 | 2,504 | 6,450,364 |
| 1M | 1000 Genomes Phase 3 chr22 | 2,504 | 1,103,548 |
| 500K | 1000 Genomes Phase 1 chr 22 | 1,092 | 490,036 |

For the synthetic dataset, we use the method provided by Wright and Ziegler [17]. The synthetic dataset consists of *n* samples and *p* features where *p >> n* and values for each feature are ordinal with three levels represented as numbers 0, 1 and 2 (which correspond to an additive effect encoding of genomic variation) randomly generated from a uniform distribution with equal probabilities. For all synthetic dataset, the response variable is a function of five randomly selected features.

The model parameters we use for simulations are 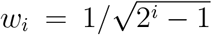 for *i* = 1,…, 5 and we set 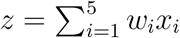. We let 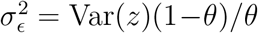 where *θ* is a parameter controlling the fraction of variance explained by the informative features, and in our study we chose *θ* = 0.125. Then *y* = *z* + *ϵ* where 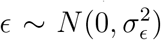. The dichotomous response is generated by thresholding *y* at the 0.5 quantile:

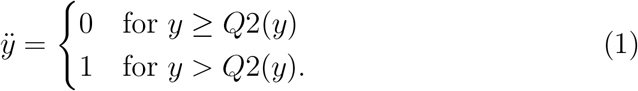

### 2.2 Computational resources

Computational platform described in Table 3 are used in our experiments.

**Table 3:** Computational resources used in our experiment

|  | Description | Number of<br>*Executors* | Total<br>Memory<br>(GB) |
| --- | --- | --- | --- |
| C1 | A *Spark* 1.6.1 cluster with 12 worker nodes each with 16 Intel Xeon E5-2660@2.20GHz CPU cores and 128 GB of memory | 32 | 32×8 |
| C2 | A *Spark* 1.6.1 cluster with 12 worker nodes each with 16 Intel Xeon E5-2660@2.20GHz CPU cores and 128 GB of memory | 130 | 130×8 |
| C3 | An Amazon aws emr cluster with r4.2xlarge machine as master and 8 r4.4xlarge machines as worker nodes | 8×16 | 8×61 |

### 2.3 *CursedForest* implementation

*CursedForest* implements the original algorithm of Breiman [5]. Given the dataset of *n* samples and *p* features, in our implementation, the dataset is partitioned by features and each partition is allocation in an *Worker* such that the *i^th^ Worker* holds *p*_*i*_ features for all samples where 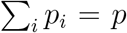. To build each tree, the *Driver* first bootstrap samples thus different bootstrapped set of samples are used to build each tree. Then, for each node of each tree *mtry* different features are randomly picked from the dataset to compute *Gini Impurity* (parallelized over all *Executors*). Since features are partitioned across *Workers*, Each *Executor* (assume one per *Worker*) randomly pick *mtry × p_i_* features and finds the best split locally for the node. The *Driver* aggregates all local best split for each node finds the global best split and updates all *Executors* about the global best split. This process is carried out across *rbs* trees in parallel where *rbs* is set by the user. Hence multiple nodes of multiple trees are processed in parallel.

We keep track of the change in *Gini Impurity* scores after splitting at each node in each tree. This information is used to calculate the *Importance Score* which is used as a metric to quantify the contribution of each feature in classifying the samples.

This implementation avoids communication bottlenecks between the *Driver* and the *Executors* as information exchange is minimal allowing it to build large numbers of trees efficiently. Furthermore, *CursedForest* has memory efficient representation of genomics data, optimized communication patterns, and computation batching. It also provides an efficient implementation of Out-Of-Bag (OOB) error calculation, which substantially simplifies parameter tuning over the more computationally expensive alternative of cross-validation.

*CursedForest* is available as a *Github* repository

The GitHub repository (https://github.com/aehrc/VariantSpark) holds information for setting up on any local or cloud-based computing environment supporting Apache *Spark*, such as *Amazon Web Services* and *Google Cloud Platform*. In addition, *CursedForest* can also be accessed through a notebook interface hosted at *Databricks* (https://aehrc.github.io/VariantSpark/notebook-examples/VariantSpark_HipsterIndex.html). Note that, in addition to *VCF* and *CSV* file format, *CursedForest* also works with the *ADAM16* data schema, implemented on top of *Avro* and *Parquet*, as well as the *HAIL API* (https://github.com/hail-is/hail) for variant pre-processing.

### 2.4 Usage of the*Gini Impurity* score as a data splitting criterion

The current implementation of *CursedForest* uses a *Gini Impurity* criterion to choose the feature for splitting at each node. This was introduced for *Decision Tree* in Breiman *et al.* [10]. For each node *A* in each tree, except leaf nodes, the program finds the feature (between randomly selected *mtry* features) that best splits samples in *A* into two child node *L* and *R*. Assume node *A* includes *m* samples each of which labelled with *q*, where *q* ∈ 1,…, *Q*. Given *f*_*q*_ as the fraction of samples in the *R* which are labeled as *q*, then the *Gini Impurity* of the node *A* (*Gini*_*R*_)is computed as,

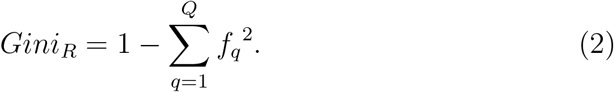

Assume *l* and *r* are the fractions of samples in *A* which are moved to *L* and *R* respectively (*l* + *r* = 1) after splitting node *A* with feature *t*. The *Information Gained* by feature *t* (*IG*_*t*_) at node *A* is computed as *Gini Impurity* of *A* minus weighted average of *Gini Impurity* of *L* and *R* using equation,

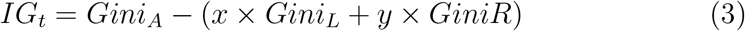

The feature that results in the highest *Information Gained* is selected as the local best split for the node by *Executor*. When the *Driver* collects all local best splits it looks for the one that maximizes *Information Gained* and chooses it as the best global split.

Feature *t* might be selected as the best split for multiple nodes of multiple trees. The raw *Importance Score* of feature *t* is defined as the mean of *IG*_*t*_ across all node in the *Random Forest* where *t* is chosen as best split.

### 2.5 Out-of-Bag training and testing procedure

The *Random Forest* algorithm builds each tree on a subset of individuals (approximately two thirds), leaving the other third out. The algorithm then tests each tree against the held-out samples, giving an estimate of the error for each tree. Averaging the error for each tree returns the OOB error for the model. According to Breiman [10], the out-of-bag estimate is as accurate as using a test set of the same size as the training set.

## 3 Results and discussion

### 3.1 *CursedForest* outperforms existing methods for multi-class classification

Figure 1a shows the execution time of all methods on different sized datasets. Note that not all programs were able to process all datasets due to out of memory errors and time-out. *CursedForest* is faster than all other methods for all dataset except for the smallest dataset (500K) where *ranger R* implementation is faster (70 seconds *vs.* 110 seconds). The *ranger R* is the second fastest implementation of *Random Forest* but cannot process the largest dataset (80M features). *CursedForest* is 4.5, 5.3 and 4.2 times faster than *ranger R* processing 1M, 6M, and 20M dataset respectively. *Google’s PLANET* is the only other *Spark* implementation of *Random Forest* but cannot process dataset larger than 6M features. *CursedForest* is 9.3 and 26.4 times faster than *Google’s PLANET* processing 1M and 6M dataset respectively.

**Figure 1:**
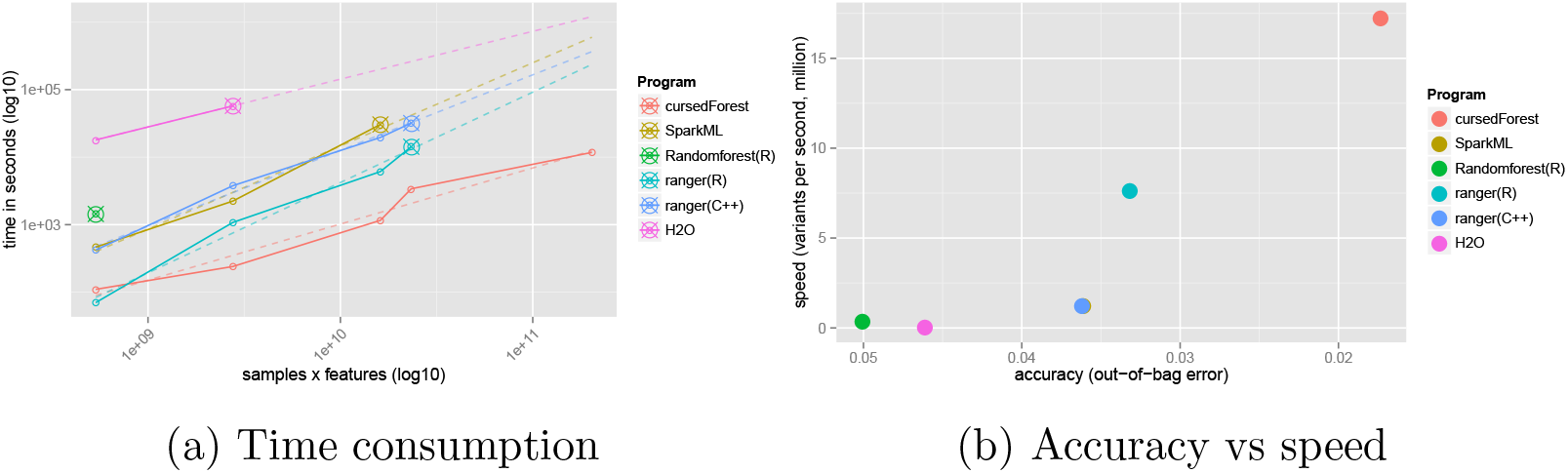
Comparison of speed and accuracy of *CursedForest* with other published methods. SparkML refers to *Google’s PLANET*. Crossed cicles (⊗) mark the last successfully process dataset by the respective methods.

*CursedForest* is the only method able to scale to the largest dataset (80M features). This demonstrates that although *Google’s PLANET* was designed to handle many samples, it is unable to efficiently cope with a large number of features. Extrapolating from this, the dotted line in Figure 1a shows that all current implementations would require between 27 hours and 11 days to finish analyzing the largest dataset (80M), while *CursedForest* completes the task in about 3 hours on a small cluster set-up (C1).

Figure 1b compares accuracy and speed of the different *Random Forest* implementations on the largest dataset they successfully complete. The accuracy is measured in terms of out-of-bag error rate (lower is better) and the speed is computed as million variants per second (higher is better). *CursedForest* is the fastest implementation and delivers the most accurate result as it was the only method able to utilize the whole genome (80M).

The runtime of *CursedForest* can be substantially improved due to its ability to efficiently utilize large numbers of commodity computers. Table 4 shows the runtime when utilizing a large *Spark* cluster (C2) (see Section 2.2) to run the same analysis as above. We can observe that the speed-up improvement grow with data set size, with up to 5-fold speed-up for 80M (from 11, 760 to 2, 214 seconds). Also, noteworthy is the reduction in error when the whole data set is utilized.

### 3.2 *CursedForestis* linearly scalable with samples, features and CPUs

Here, we explore the performance of *CursedForest* in more detail by testing its ability to scale beyond the real dataset size. We hence generate synthetic datasets with up to 50 million features (i.e., *p* = 50,000,000), with up to 10,000 samples (see Section 2.1).

We measured the runtime to build a binary *Random Forest* model of 100 trees with a fixed *mtry* fraction of 0.25 using our C2 computer (see Section 2.2) with these different synthetic datasets. *CursedForest* scales linearly with increase in feature sizes *p* and sublinearly with the increase in sample size *n* (see Figure 2a). *CursedForest* also scales well when increasing the number of CPUs making cloud-application with on-demand cluster sizes possible (see Figure 2b). As shown in Table 4 the 80M dataset can be processed in about 37 minutes. With increase in the number of executors the execution time can further decreases.

**Figure 2:**
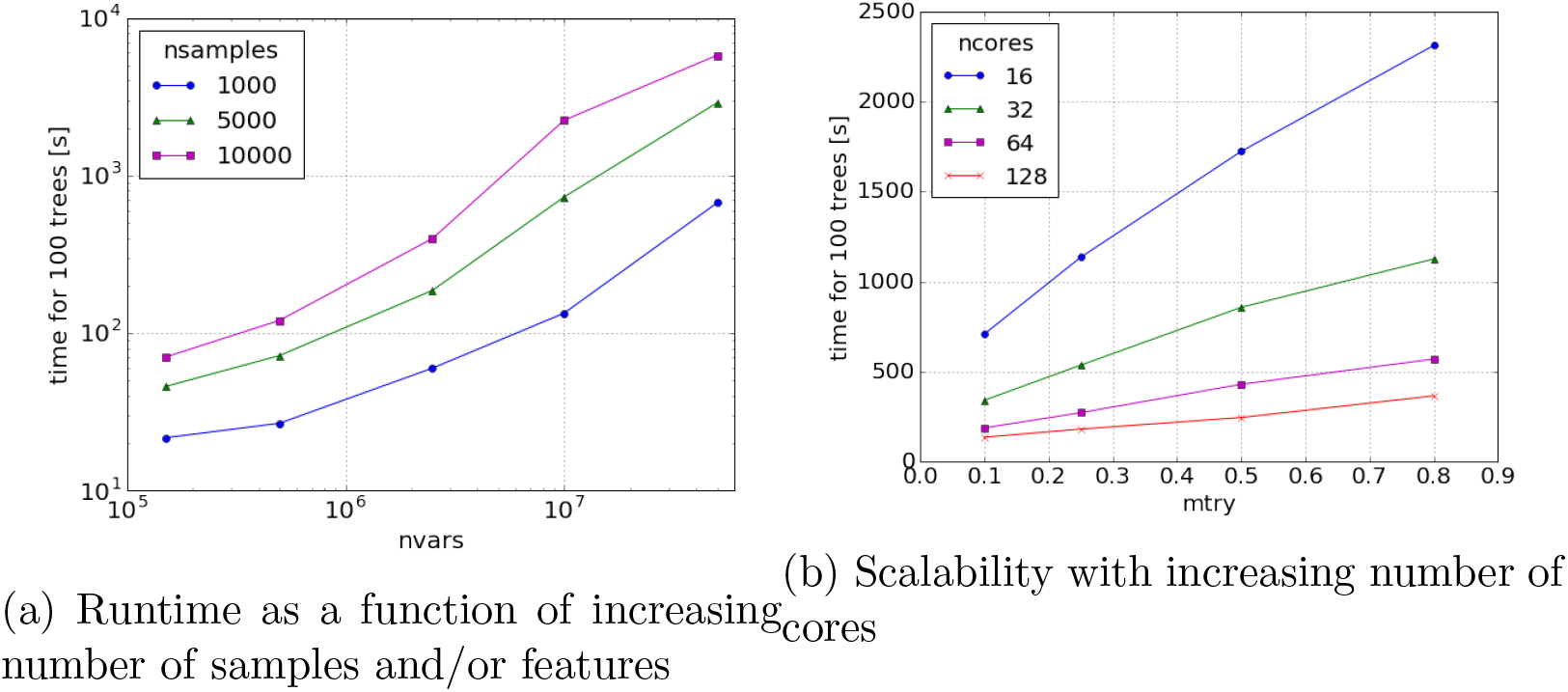
Performance in response to varying *ntree* and *mtry*

**Table 4:** Accuracy and scalability of a multi-class classification with 50 trees using *CursedForest*

| Dataset | Error Rate<br>(OOB) | Runtime (min) |  | Speed up |
| --- | --- | --- | --- | --- |
|  |  | C2 | C1 |  |
| 1M | 0.06 | 1.6 | 4.0 | 2.5 |
| 6M | 0.04 | 4.2 | 19.3 | 4.6 |
| 20M | 0.02 | 10.8 | 56.7 | 5.2 |
| 80M | 0.01 | 36.9 | 196.0 | 5.3 |

### 3.3 Wide data and the choice of *mtry* and *ntree*

*CursedForest* is a partial implementation of the original algorithm of [5]. As such the choice of *ntree* and *mtry* follows the same logic. However, the original choices of the default values of these parameters was based on Breiman’s experience with a number of data sets [5]. *CursedForest* may be operating in regions where a different set of heuristics may be needed to guide the parameter settings.

In this section, we test the limits of *CursedForest* in association and classification analysis on high-dimensional data. For this purpose, we generated a synthetic dataset with 2.5 million features (i.e., *p* = 2,500,000), of which 5 are designed to be related to the response variable, with 5,000 samples (i.e. *n* = 5,000) (see Section 2.1). We use our C2 computer (see Section 2.2) for this analysis.

We fit the *Random Forest* model and estimate the classification accuracy by capturing the OOB error. We also measure the feature selection performance by capturing the rank-biased overlap (RBO) measure [16]. RBO assesses whether *CursedForest* is able to retrieve the 5 features in order of their association weight as a scale from 0 (no feature recovered) to 1 (fully recovered).

See Figure 3a for plots of these two measures for this example. Note that, for the parameter *mtry*, the plot shows the proportion, *mtry/p*, which means that the default value of 1581 is shown as 1581/*p* ≈ 0.0006. The default value for *mtry* does not result in a good classification performance for this large feature dataset. The OOB for this value of *mtry* does not drop below 0.5, even when the number of trees is increased. Increasing *mtry* in combination with *ntree* yields the best performance with the OOB error essentially constant around 0.4 across a large range of *mtry* and *ntree* values. This is in contrast to the feature-selection performance, where the RBO measure heavily depends on *ntree* and gives better results with lower values of *mtry* (Figure 3b).

**Figure 3:**
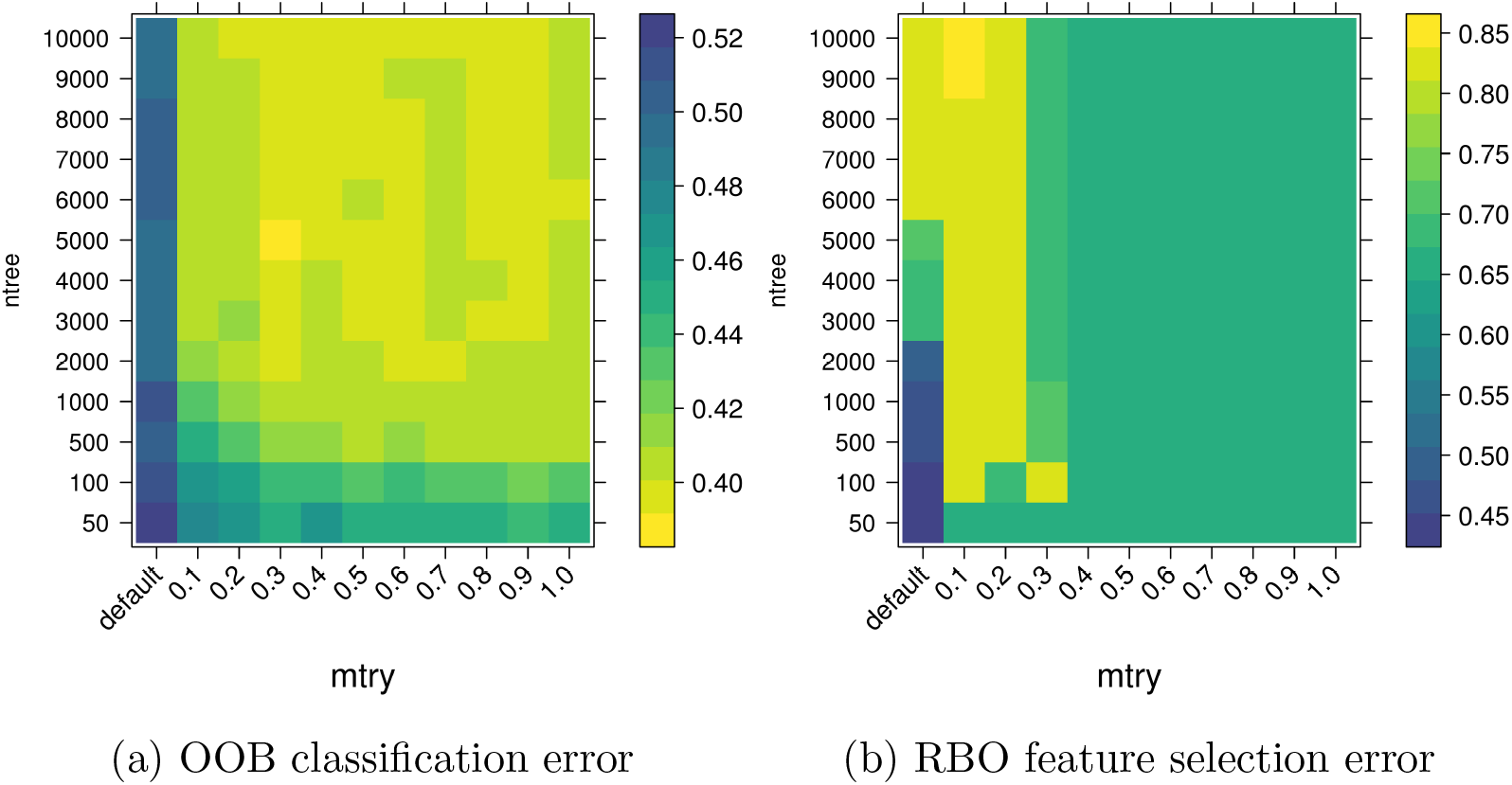
Accuracy performance in response to varying *ntree* and *mtry*. *mtry* is given as a fraction of *p*.

This may be because a large *mtry* leads to more correlated trees as the same important features have a higher chance of being selected in all trees, which does not yield good performance outcomes. This issue is less pro-nounced for classification error where random features can mimic the response variable, hence resulting more in a performance plateau. Increasing the number of trees, on the other hand, improves performance especially when the trees are kept diverse (small *mtry*) but appropriate for large feature datasets (*mtry* larger than default). The ability of *CursedForest* to build decision trees in parallel at the compute node level hence caters perfectly to the requirement of large feature datasets as more trees can be built given a fixed time budget.

### 3.4 *CursedForest* tree-building method is faster than *Yggdrasil*

Here we compare the tree building implementation of *CursedForest* to *Yggdrasil*. Both methods partition the data by features to parallelize Gini computation, which was demonstrated to yield a better performance for wide data than *Google’s PLANET*, which partitions by samples [1]. Since *CursedForest* is an ensemble method, we approximate the single *Decision Tree* functionality of *Yggdrasil* by (1) disabling bootstrapping, (2) setting *mtry* = *p*, (3) setting *nTree* = 1.

The largest dataset we were able to process with *Yggdrasil* is dataset 1M. It took 102 seconds (average over two runs) for *Yggdrasil* to build a *Decision Tree* for this dataset, compared to 3 seconds for *CursedForest* (average over two trees, *mtry* = *p* and without bootstrapping). This shows nearly 33 times speed up over *Yggdrasil*. Enabling parallel tree-building reduces the runtime for *CursedForest* to an average of 1.14 second to build each tree, which represents a 89-fold speed-up over the *Yggdrasil* implementation. The time mentioned above only include the actual training process, not data loading and other auxiliary steps. For this experiment we use C3 cluster as described in Table 3.

## 4 Conclusion

The challenge of “big” and “wide” data is especially pronounced in the biomedical space where dataset acquisition is predicted to far outpace that of traditional “Big Data” disciplines [15]. Catering for this, we extended *Random Forest* to cope with extremely high dimensional data using a novel parallelization approach enabled by *Spark*. Compared to *Google’s PLANET* and other non-*Spark* implementations, as well as the purpose-designed *Yggdrasil*, *CursedForest* can scale to millions of features. It also offers the fastest trainings method for *Random Forest* on a wide range data-sets sizes compared to the other tools tested.

## References

[1] F. Abuzaid, J. K. Bradley, F. T. Liang, A. Feng, L. Yang, M. Zaharia, and A. S. Talwalkar. Yggdrasil: An Optimized System for Training Deep Decision Trees at Scale, volume 29, pages 3817–3825. Curran Associates, Inc., 2016.

[2] D. C. Bauer, C. Gaff, M. E. Dinger, M. Caramins, F. A. Buske, M. Fenech, D. Hansen, and L. Cobiac. Genomics and personalised wholeof-life healthcare. Trends in Molecular Medicine, 20(9):479–486, 2014.

[3] B. P. Bayardo, J. S. Herbach, S. Basu, and R. J. Planet: Massively parallel learning of tree ensembles with mapreduce. In Proceedings of the 35th International Conference on Very Large Data Bases (VLDB2009), 2009.

[4] R. Bellman and R. Bellman. Adaptive Control Processes: A Guided Tour. Princeton University Press, 1961.

[5] L. Breiman. Random forests. Machine Learning, 45(1):5–32, 2001.

[6] T. Chen and C. Guestrin. Xgboost: A scalable tree boosting system. In Proceedings of the 22nd acm sigkdd international conference on knowledge discovery and data mining, pages 785–794. ACM, 2016.

[7] W. T. C. C. Consortium. Genome-wide association study of 14,000 cases of seven common diseases and 3,000 shared controls. Nature, 447(7145):661–678, 2007.

[8] H2O. Open-source machine learning platform for enterprises, https://www.h2o.ai/h2o/.

[9] L. Lello, S. G. Avery, L. Tellier, A. Vazquez, G. d. l. Campos, and S. D. Hsu. Accurate genomic prediction of human height. arXiv preprint arXiv:1709.06489, 2017.

[10] C. J. S. R. O. Leo Breiman, Jerome Friedman. Classification and Regression Trees. Wadsworth Publishing Company, Belmont, California, U.S.A., 1 edition, 1984.

[11] A. E. Locke, B. Kahali, S. I. Berndt, A. E. Justice, T. H. Pers, et al. Genetic studies of body mass index yield new insights for obesity biology. Nature, 518(7538):197–206, 2015.

[12] B. Loebbecke and A. Picot. Reflections on societal and business model transformation arising from digitization and big data analytics: A research agenda. The Journal of Strategic Information Systems, 24(3):149–157, 2015.

[13] A. R. O’Brien, N. F. W. Saunders, Y. Guo, F. A. Buske, R. J. Scott, and D. C. Bauer. Variantspark: population scale clustering of genotype information. BMC Genomics, 16(1), 2015.

[14] N. Siva. 1000 genomes project, 2008.

[15] Z. D. Stephens, S. Y. Lee, F. Faghri, R. H. Campbell, C. Zhai, M. J. Efron, R. Iyer, M. C. Schatz, S. Sinha, and G. E. Robinson. Big data: Astronomical or genomical? PLoS Biol, 13(7):e1002195, 2015.

[16] W. Webber, A. Moffat, and J. Zobel. A similarity measure for indefinite rankings. ACM Transactions on Information Systems, 28(4):20:1–20:38, 2010.

[17] M. N. Wright and A. Ziegler. Ranger: A fast implementation of random forests for high dimensional data in c++ and r. Journal of Statistical Software, 2016.

[18] M. N. Wright, A. Ziegler, and I. R. König. Do little interactions get lost in dark random forests? BMC Bioinformatics, 17(1):145, 2016.

